# Functional characterization of peroxisome biogenic proteins Pex5 and Pex7 of *Drosophila*

**DOI:** 10.1101/366633

**Authors:** Francesca Di Cara, Richard A. Rachubinski, Andrew J. Simmonds

## Abstract

Peroxisomes are ubiquitous membrane-enclosed organelles involved in lipid processing and reactive oxygen detoxification. Mutations in human peroxisome biogenesis genes (*Peroxin*, *PEX*) cause progressive developmental disabilities and, in severe cases, early death. PEX5 and PEX7 are receptors that recognize different peroxisomal targeting signals called PTS1 and PTS2, respectively, and traffic proteins to the peroxisomal matrix. We characterized mutants of *Drosophila melanogaster Pex5* and *Pex7* and found that adult animals are affected in lipid processing. Moreover, *Pex5* mutants exhibited severe developmental defects in the embryonic nervous system and muscle, similar to what is observed in humans with *Pex5* mutations, while *Pex7* fly mutants were weakly affected in brain development, suggesting different roles for Pex7 in fly and human. Of note, although no PTS2-containing protein has been identified in *Drosophila*, Pex7 from *Drosophila* can function as a *bona fide* PTS2 receptor because it can rescue targeting of the PTS2-containing protein Thiolase to peroxisomes in *PEX7* mutant human fibroblasts.

## INTRODUCTION

Peroxisomes have long been known to be involved in a variety of important biochemical functions, most notably lipid metabolism and the detoxification of reactive species (Bowers, 1998; Deduve, 1965; Nguyen et al., 2008; Wanders and Waterham, 2006). Recent evidence has shown peroxisomes to have important roles in development, immune signaling and viral maturation. Peroxisome biogenesis genes (*Peroxin*, *PEX*) are for the most part conserved across the breadth of eukaryotes (Platta and Erdmann, 2007; Schrader and Fahimi, 2006). Thirteen *PEX* genes are required for peroxisome biogenesis in humans, and mutations in these genes cause the peroxisome biogenesis disorders (PBDs), which manifest as heterogeneous syndromes with varied developmental defects (Braverman et al., 2013). PEX5 and PEX7 act as receptors that recognize *cis*-acting signals called peroxisome targeting signals (PTS) in soluble peroxisomal proteins to traffic them from the cytosol to the peroxisome matrix (Smith and Aitchison, 2013) (Ito et al., 2007; Klein et al., 2001; Purdue et al., 1997). PEX5 and PEX7 homologues are found across the eukaryota, including kinetoplastids, yeasts, plants and mammals (Kanzawa et al., 2012; Kragler et al., 1998; Lazarow, 2006; Matsumura et al., 2000; McCollum et al., 1993; Purdue et al., 1997; Rehling et al., 1996; Woodward and Bartel, 2005). PEX5 recognizes the C-terminal PTS1 with the canonical sequence Ser-Lys-Leu (SKL), while PEX7 recognizes an N-terminal nonapeptide PTS2 with the consensus sequence (R/K)(L/V/I)X5(H/Q)(L/A) (Ito et al., 2007; McCollum et al., 1993; Rehling et al., 1996; Shimozawa et al., 1999). Mutation of *PEX5* gives rise to Zellweger syndrome, while mutation of *PEX7* gives rise to Rhizomelic Chondrodysplasia Punctata Type 1 (RCDP1) (Purdue et al., 1997).

Mutation of *Drosophila Pex* genes is linked to a range of phenotypes, including lethality (*Pex1*, *Pex3*, *Pex19*) and male sterility (*Pex16*) (Beard and Holtzman, 1987; Bulow et al., 2018; Chen et al., 2010; Faust et al., 2014; Mast et al., 2011; Nakayama et al., 2011). In *Drosophila* S2 cells, knock down of *Pex5* transcript reduces targeting of PTS1-containing proteins to peroxisomes function, while depletion or overexpression of *Pex7* transcript led to smaller or larger peroxisomes, respectively, than normal (Baron et al., 2016). However, the actual function of *Drosophila* Pex7 remains unclear, as no *bona fide* peroxisomal PTS2-containing protein has been identified in *Drosophila*; fly homologues of peroxisomal proteins trafficked by the PTS2/PEX7 import pathway, *e.g*. peroxisomal Thiolase, use the PTS1/Pex5 pathway in *Drosophila* (Baron et al., 2016; Faust et al., 2012).

Here we show that *Drosophila Pex5* mutants exhibited severe developmental defects in the embryonic nervous system and muscle, similar to what is observed in Zellweger syndrome patients with *PEX5* mutations. *Pex7* fly mutants exhibited minor defects in brain development. We also show that *Drosophila* Pex7 can function as a *bona fide* PTS2 receptor because it can rescue targeting of the PTS2-containing protein Thiolase to peroxisomes in *PEX7* mutant human fibroblasts.

## RESULTS AND DISCUSSION

### *Drosophila Pex5* is required for development

The *Pex5^MI06050^* mutation was caused by a MiMIC insertion disrupting the coding region (Venken et al., 2011)*. Pex5* is on the X chromosome and *Pex5^MI06050^* mutants were lethal when homozygous or as hemizygous males with only 20% of embryos hatching (Figure 1A-C). A further 15% of *Pex5^MI06050^*/*Pex5^MI06050^* mutants died as larvae with only 5% pupating (Figures 1A1-3, C). The 2% of pupae survived died at eclosion (Figure 1A1-4, 1C). *Pex5^MI06050^*/*Pex5^MI06050^* embryos had 35% of *Pex5* mRNA compared to control (Figure 1B). Maternally provided *Pex5* mRNA likely caused the phenotypic variability observed. Finally, strains where the *Pex5^MI06050^* MiMIC element was excised precisely were viable supporting that the phenotypes were due to *Pex5* disruption.

**Fig. 1.**
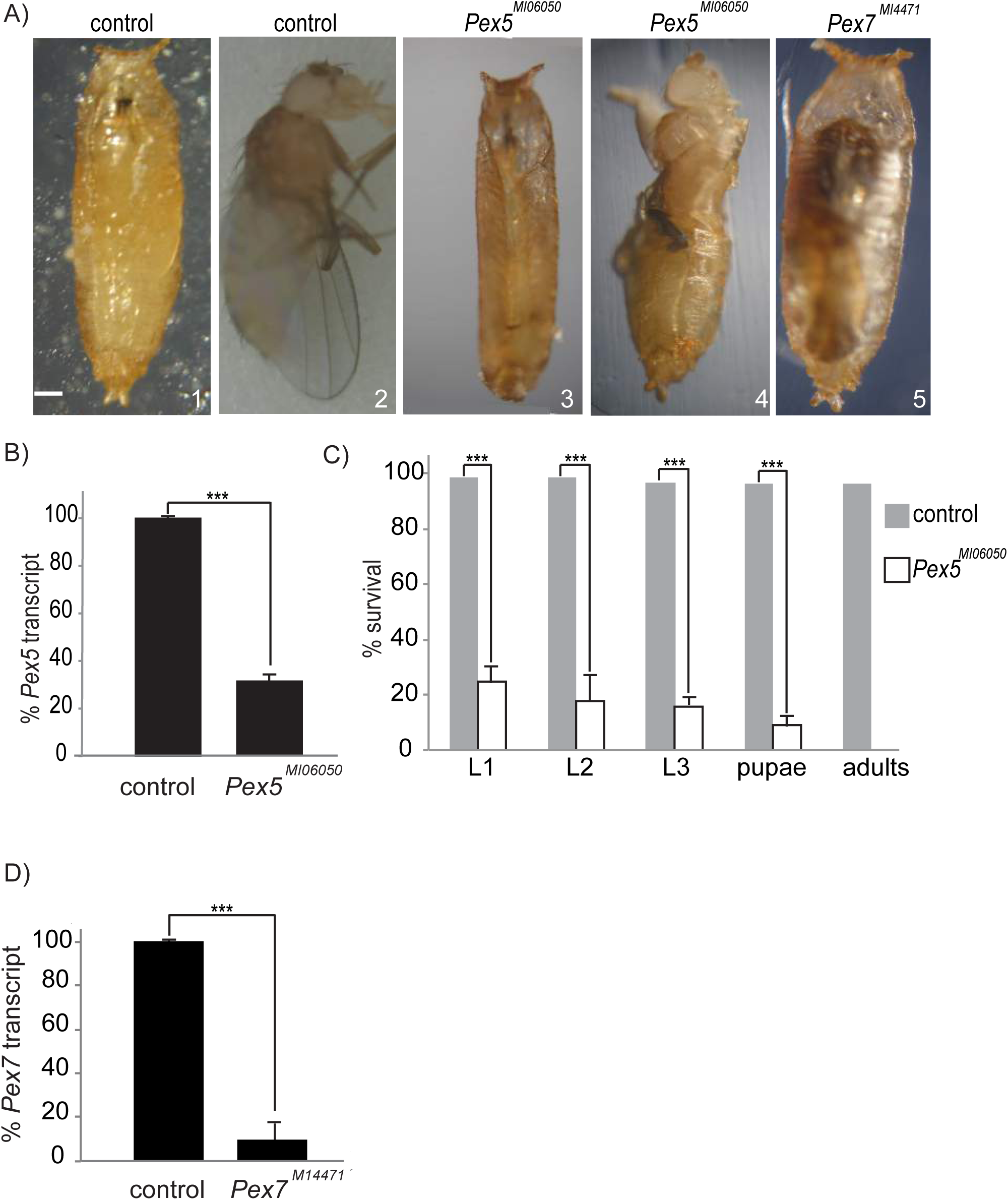
*Pex5* and *Pex7* mutants present developmental defects. (A) 1. Control pupa 7 days after egg laying (AEL). 2. Control adults eclosed at day 9 AEL. 3. Most *Pex5^MI06050^*/*Pex5^MI06050^* mutants arrest at pseudo-pupa stage (day 11 AEL) 4. Some *Pex5^MI06050^*/*Pex5^MI06050^* mutants die during eclosion (day 10 AEL). 5. Some *Pex7^MI4471^*/*Pex7^MI4471^* mutants arrest at pupa stage (day 11 AEL). Scale bar=1mm. (B) QRTPCR confirms reduction in the amount of *Pex5* mRNA in *Pex5^MI06050^*/*Pex5^MI06050^* embryos, relative to controls. Values are averages of 4 independent experiments ± SD. Significance was determined using Student’s *t*-test; *** p < 0.001. (C) Most *Pex5^MI06050^*/*Pex5^MI06050^* mutants die at embryo stage and none eclose as adults. Values are averages of 4 independent experiments ± SD. N=200 embryos were analyzed for each genotype. Significance was determined using Student’s *t*-test; *** p < 0.001. (D) QRTPCR confirms reduction of *Pex7* mRNA in *Pex7^MI4471^*/*Pex7^MI4471^* embryos, relative to control. Values are averages of 4 independent experiments ± SD. Significance was determined using Student’s *t*-test; *** p < 0.001.

Human *PEX5* mutations cause central nervous system (CNS), peripheral nervous system (PNS) and musculature defects (Braverman et al., 2013; Steinberg et al., 2006). Thus, we assayed CNS and PNS organization in *Pex5^MI06050^* mutant embryos compared to age-matched controls. In *Pex5^MI06050^*/*Pex5^MI06050^* embryos, both the PNS and ventral nerve cord (VNC) were disorganized (Figure 2A). PDB patients often show axonal demyelination (Braverman et al., 2013) and while *Drosophila* neurons are unmyelinated, wrapping glia play an analogous role (Freeman and Doherty, 2006; Matzat et al., 2015). Glial cells were disorganized in *Pex5^MI06050^*/*Pex5^MI06050^* embryos compared to controls (Figure 2A). Finally, the developing longitudinal and oblique musculature in late-stage *Pex5^MI06050^*/*Pex5^MI06050^* embryos was also disorganized (Figure 2B).

**Fig. 2.**
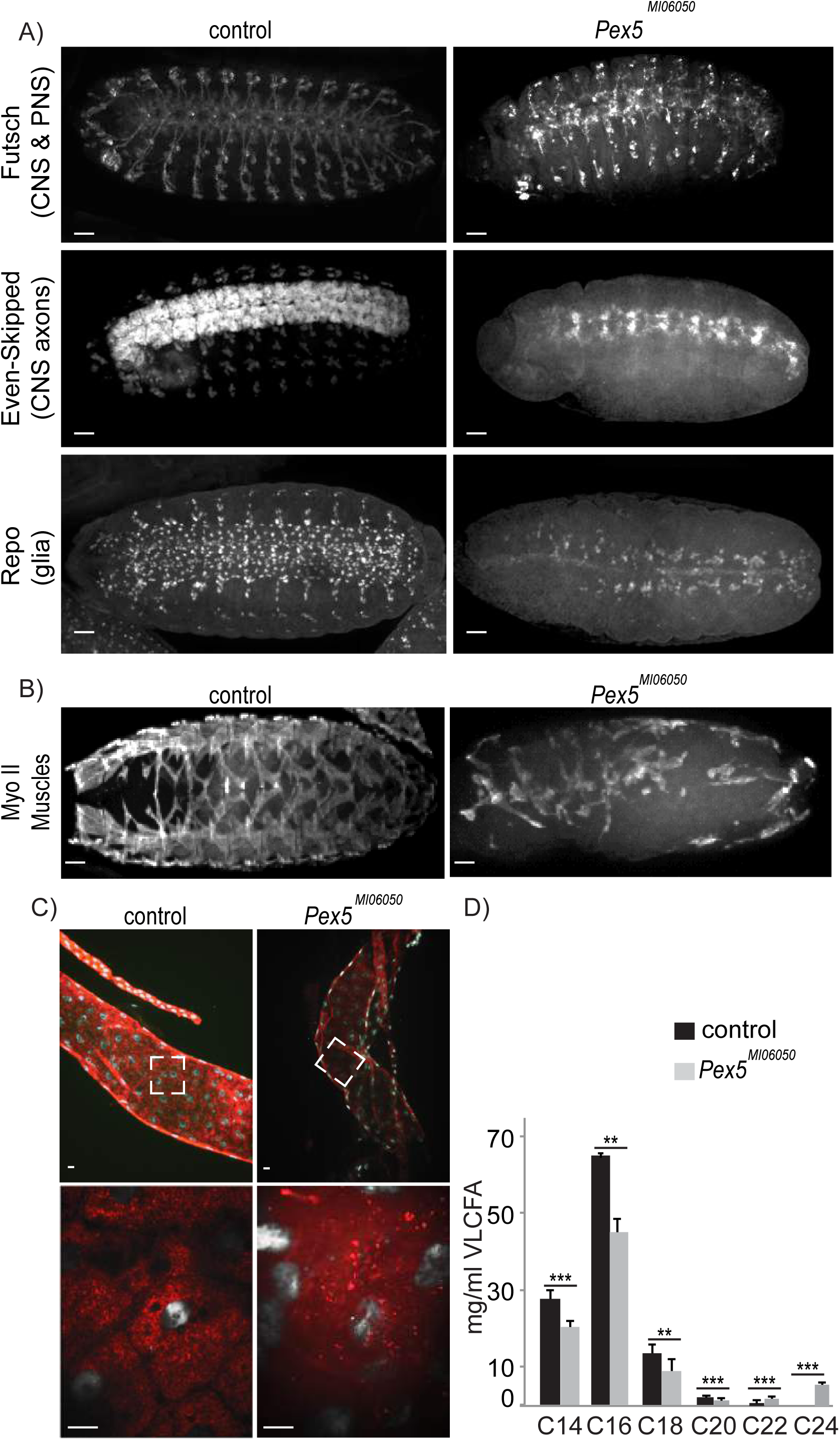
The *Pex5^MI06050^* mutation affects NS, PNS and muscle development. (A) The repeated segmental pattern of neuron of the CNS and PNS (marked by anti-Futsch), CNS axons (marked by anti-Even-skipped) and glia cells (except midline glia, marked by anti-Repo) in control embryos (stage 15). This pattern is severely disrupted in *Pex5^MI06050^*/*Pex5^MI06050^* embryos. Scale bar=10μm. (B) The repeated segmental pattern of developing muscles marked by Anti-Myosin II in control embryos is disrupted in *Pex5^MI06050^*/*Pex5^MI06050^* embryos (stage 15). Scale bar=10μm. (C) Mature peroxisomes (punctate anti-SKL signal, red) are observed in the larval midgut. In *Pex5^MI06050^*/*Pex5^MI060^* mutants, diffuse SKL aggregates indicate PTS1 import was impaired. DAPI labelled nuclei are in grey. The boxed region in the top panel is shown magnified in the lower panel. Scale bar=10 µm (D) *Pex5^MI06050^*/*Pex5^MI060^* mutants have lower long chain fatty acid levels (C_14_, C_16_ and C_18_) and accumulate VLCFAs (C20, C22 and C24) compared to controls. Values are averages of 4 independent experiments ± SD. N=1000 larvae per sample per each genotype were used in each replicate (4000 total). Significance was determined using Student’s *t*-test; *** p < 0.001; ** p<0.01.

### *Drosophila Pex5* is required for peroxisome biogenesis

To determine if the *Pex5^MI06050^* mutation affects PTS1-mediated peroxisome import we analyzed the localization of PTS1/SKL containing proteins in third instar larvae midgut cells (Szilard et al., 1995). A punctate signal, corresponding to peroxisomes with active PTS1 import, was present only in controls. Large SKL-positive aggregates were observed in *Pex5^MI06050^*/*Pex5^MI06005^* midgut cells indicating lack of PTS1 import (Figure 2C). To examine the systemic effect of *Pex5^MI06050^* mutation on peroxisome function, we profiled the spectrum of fatty acids. *Pex5^MI06050^*/*Pex5^MI06050^* embryos accumulate VLCFAs (C22 and C24) and relatively lower levels of C14, C16, C18 and C20 compared to controls (Figure 2D).

### *Drosophila Pex7* is required for neuronal development

*Pex7^MI4471^* is caused by MiMIC insertion into the *Pex7* coding region (Venken et al., 2011)*. Pex7^MI4471^*/*Pex7^MI4471^* Unlike *Pex5^MI06050^*, mutants were viable with only 5% arresting before pupal stage. Arrested pupae were the same size as control but developmental abnormalities were observed (Figure 1A1, 2, 5). QRTPCR analysis showed *Pex7^MI4471^*/*Pex7^MI4471^* embryos had only 10% *Pex7* mRNA relative to controls (Figure 1D). As a symptom of *PEX7*-linked RDCP is defects neurogenesis (Braverman et al., 2013; Steinberg et al., 2006), we analyzed the brain morphology of *Pex7^MI4471^*/*Pex7^MI4471^* third instar larvae. Compared to wild type, *Pex7^MI4471^*/*Pex7^MI4471^* larvae were larger (Figure S1A), but their brain volume was smaller (Figure 3A). Although *Pex7^MI4471^*/*Pex7^MI4471^* animals were slightly larger than control at the same stage but the mutation did not appear to affect the developmental timing as most of the animals reached the adult stage at the same time as control animals (Figure 1A1,2, 5). This phenotype unusual in that small brain / enlarged body size is usually associated with developmental arrest (Colombani et al., 2005; Mirth et al., 2005).

**Fig. 3.**
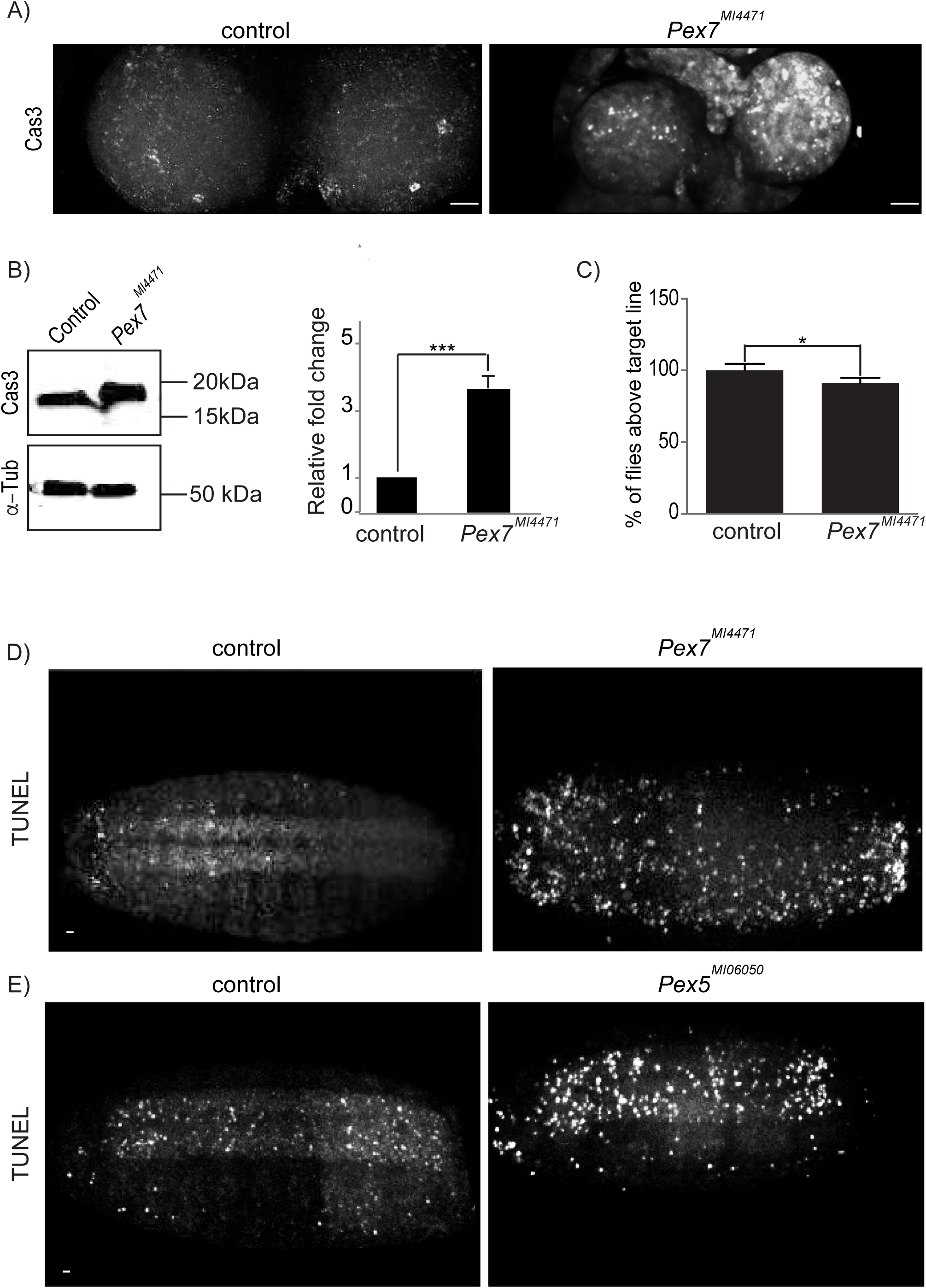
*Pex7* mutation causes defects in CNS development. (A) *Pex7^MI4471^*/*Pex7^MI4471^* larvae that arrest between second and third instar had smaller brains than control. The number of apoptotic cells (marked by activated Caspase3, CM1) was higher in *Pex7^MI4471^*/*Pex7^MI447^* brains. Scale bar=10 µm. (B) Total CM1 levels were higher *Pex7^MI4471^*/*Pex7^MI447^* second and third instar larval brains. Values represent averages of 4 independent experiments ± SD. Significance was determined using Student’s *t*-test; *** p < 0.001. (C) *Pex7^MI4471^*/*Pex7^MI4471^* show reduced performance in a climbing assay, testing coordinated locomotion, than control. Values represent averages of 12 independent experiments ± SD. Significance was determined using Student’s *t*-test; * p < 0.05. (D) TUNEL positive cells were detected around the embryonic CNS in *Pex7^MI4471^*/*Pex7^MI447^* embryos compared to control. (E) *Pex5^MI06050^*/*Pex5^MI06050^* embryos also had elevated numbers of TUNEL postive cells in the CNS. Images are representative of 4 independent experiments. N=20 per experiment/genotype. Scale bar=10µm.

Reduced brain size could be due either to reduced proliferation or excess cell death (apoptosis) (Shklyar et al., 2014). The number of phospho-Ser10-histone 3 (PH3) marked mitotic cells (Wei et al.; 1999) was similar in *Pex7^MI4471^*/*Pex7^MI4471^* brains and controls (Figure S1B). *Pex7^MI4471^*/*Pex7^MI4471^* brain extracts had elevated Active Caspase3 (CM1) levels compared to control (Figure 3B). Also, there were more CM1 positive cells within developing *Pex7^MI4471^*/*Pex7^MI4471^* brains compared to controls (Figure 3A). Accumulation of TUNEL positive cells in the developing brain was also seen in *Pex7^MI4471^*/*Pex7^MI4471^* and *Pex5^MI06050^*/*Pex5^MI06050^* embryos (Figure 3D, E).

Despite having altered brain morphology, *Pex7^MI4471^*/*Pex7^MI4471^* mutants are viable. Thus, we functionally assayed neural and muscular function by analyzing the ability of adult flies to exhibit a negative geotaxis response (Madabattula et al., 2015). This climbing assay is used commonly to assay the effects of CNS neurodegeneration in flies (Feany and Bender, 2000). Control flies were more successful in the climbing assay that *Pex7^MI4471^*/*Pex7^MI4471^* flies (Figure S1B). Notably, even at the end of the assay (120sec), many *Pex7^MI4471^*/*Pex7^MI4471^* flies had not completed the task (Figure 3C, S1B).

### *Drosophila* Pex7 has a role in peroxisome fatty acid processing

Lack of *Drosophila* PTS2 import calls into question if *Pex7^MI4471^*/*Pex7^MI4471^* associated defects are linked to peroxisome dysfunction. Non-esterified fatty acids (NEFAs) accumulate in flies with impaired peroxisomes (Bulow et al., 2018). Thus, we measured the relative concentration of circulating NEFAs and observed accumulation in *Pex7^MI4471^*/*Pex7^MI4471^* larvae compared to control (Figure S1D).

### *Drosophila* Pex7 can functionally substitute for human PEX7

In human cells, Thiolase is targeted to peroxisomes via PTS2/PEX7 (Braverman et al., 1997). To determine if *Drosophila* Pex7 retained a similar functional activity, we assayed Thiolase import in human fibroblasts from a RCDP patient with a mutation in *PEX7* (Braverman et al., 1997; Purdue et al., 1997). Transfection with a human PEX7 cDNA restored Thiolase import to peroxisomes (Figure 4, S3). Transfection with a *Drosophila Pex7* cDNA also rescued Thiolase import indicating *Drosophila* Pex7 is competent to mediate PTS2 import (Figure 4, S3).

**Fig. 4.**
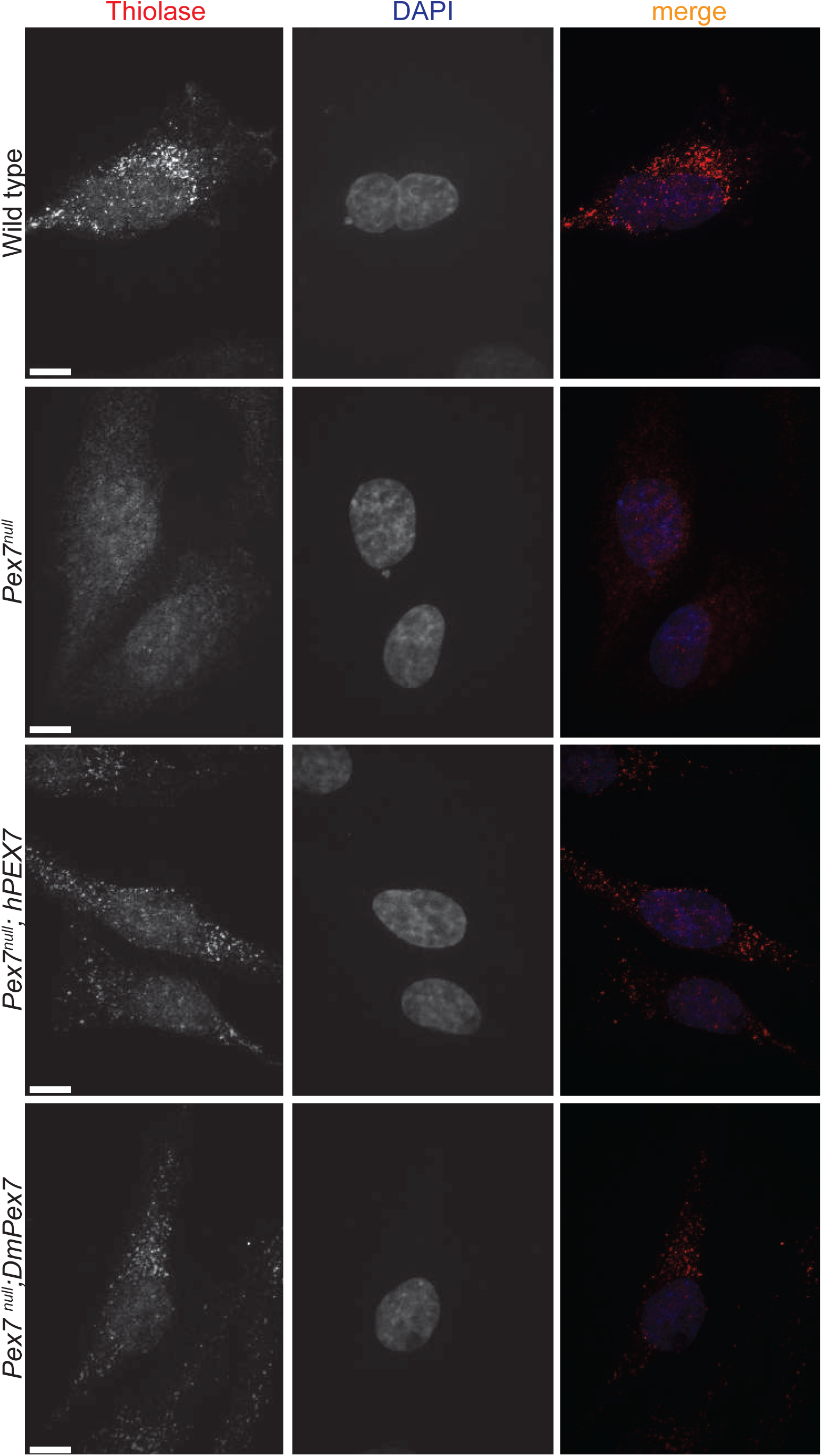
Drosophila Pex7 can facilitate PTS2 import in human cells. In wild type human fibroblasts, anti-Thiolase marks punctate cytoplasmic spots co-localizing with a peroxisome membrane marker antiPMP70. *PEX7* mutant fibroblasts (*PEX7*^null^) do not have punctate Thiolase signal indicating lack of import but PMP70 puncta remain, indicating peroxisomes are present. Transfection of *PEX7*^null^ cells with *PEX7 cDNA* restores Thiolase import to peroxisomes. Expression of a *Drosophila Pex7 cDNA* in *PEX7 null* human fibroblasts also restored Thiolase import to peroxisomes. Scale bar=10µm.

### The role of *Drosophila* Pex7 is divergent

Given the previously observed localization of Pex7 to peroxisomes, and the effect on peroxisome size (Baron et al., 2016; Faust et al., 2012; Mast et al., 2011), and effects on lipid processing and brain development we observe in *Pex7^MI4471^*/*Pex7^MI4471^* mutants, *Pex7* clearly has a role in peroxisome biogenesis or function. The lack of a canonical PTS2 trafficking pathway in flies, calls into question the mode of action. *Drosophila* Pex7 homologues of known yeast and human PTS2 targeted proteins have a PTS1 signal (Baron et al., 2016; Faust et al., 2012). It is likely that the canonical PTS2 pathway is not active in *Drosophila*, as S2 cells cannot import a canonical PTS2-mCherry reporter into peroxisomes while mammalian cells can (Faust et al., 2012). On the other hand, comparison of the amino acid sequence of *Drosophila melanogaster Pex7* to *Saccharomyces cerevisiae*, *Arabidopsis thaliana*, *Danio rerio* and *Homo sapiens* homologs (Figure S2) suggests conservation. *Caenorhabditis elegans* does not have a PTS2 pathway but does not have a *Pex7* homologue either (Motley et al., 2000). It is possible *Drosophila* has divergent PTS2 signal. Thus, to determine *Drosophila* does have a divergent PTS2 signal or Pex7 plays a PTS2-independent peroxisomal function, additional studies are needed.

## MATERIALS AND METHODS

### Cell culture

Human fibroblasts were cultured in Dulbecco’s modified Eagle’s medium (ThermoFisher) supplemented with 10% (FBS), 50U penicillin/ml and 50μg streptomycin sulfate/ml.

Fly husbandry, egg collection and survival assays

*y^1^Mi{y^+mDint2^=MIC}Pex5^MI06050^ w^*^/FM7h*) and *y^1^ w*^*^; *Mi{y^+mDint2^=MIC}Pex7^MI14471^* mutant lines and FM7(GFP) Df(1)JA27/*FM7c*, P{*w*^+mC^=GAL4-*Kr*.C}DC1, P{*w*^+mC^=UAS-GFP.S65T}DC5, *sn*^+^ strains were obtained from Bloomington Drosophila Stock Center (BDSC). *w^1118^* was used as a control in all experiments. *Drosophila* were maintained at 25°C on standard BDSC cornmeal medium. *Pex5^MI06050^* mutants balanced over FM7(GFP) were allowed to lay eggs on apple juice agar plates for two days. On the third day, embryos were collected every 2h. GFP-negative embryos were incubated on apple juice agar plates at 25°C. After 24h hatched larvae were transferred to standard cornmeal medium and surviving animals were counted at the same time each day.

### Geotaxis (climbing) assay

This assay was performed as described previously (Madabattula et al., 2015) using 20 flies (7 days old) and a 250mL glass graduated cylinder (ThermoFisher), sealed with wax film to prevent escape. Assays were conducted in ambient light at 22°C and performed at the same time each day.

### Lipid analysis

One thousand first instar (L1) larvae (equivalent to 1 mg of protein extract) were homogenized in 1ml PBS buffer and sonicated for 5min (BioRuptor) at low power. Lipids were extracted using chloroform:MeOH, 2:1 as described previously (Folch et al., 1957). 5µg of C17 dissolved in chloroform was used as an internal control. Isolates were centrifuged at 3400g and the chloroform phase containing the lipid fraction passed through a sodium sulfate column (GE Bioscience). The eluate was dried under inert gas (N_2_) and resuspended in 100µl µl of HPLC-grade hexane. 10µl was injected into an Agilent 6890 Gas Chromatograph with Flame-Ionization Detector. VLCFA concentration was normalized to the relative amount of protein determined using a Qubit II fluorimeter (ThermoFisher). Non-esterified fatty acids were analysed as per (Bulow et al., 2018).

### QRTPCR analysis

Samples were rinsed twice with PBS, and total RNA extracted using the RNeasy-Micro Kit (Qiagen). 0.5–1µg of RNA was reverse-transcribed using an iScript cDNA Synthesis kit (Biorad), QPCR was performed (Realplex, Eppendorf) using KAPASYBR Green PCR master mix (KAPA Biosystems). Samples were normalized to *RpL23* based on the ΔΔCT method. A Student’s t-test was used to calculate significance of differences in gene expression between averaged sample pairs. QRTCPR primer sequences used were: *RpL23*, 5’-GACAACACCGGAGCCAAGAACC, 5’-GTTTGCGCTGCCGAATAACCAC *Pex5*, 5’-AAATGCGAAGACATGGAACC, 5’-TGTAACGCACACGGATGAAG *Pex7*, 5’-TCGAAATAGCCAGGCCATCAAG, 5’AAGGAACCGAAGACAAGGACTC All QRTPCR data shown here are based on 3 biological samples each tested in triplicate.

### Protein analysis

50µl of cold Ephrussi–Beadle Ringer’s solution supplemented with 10mM EDTA, 10mM DTT, 1x Complete protease inhibitor and 1x PhosStop phosphatase inhibitor (Roche) was added to 3 × 10^6^ pelleted cells. 25µl of 70°C 3X SDS-PAGE Buffer (Biorad) containing 10mM DTT was added to the homogenate and incubated at 100°C for 10 min. Samples were resolved by SDS-PAGE on 10% acrylamide gels and transferred onto nitrocellulose membranes (BioRad). Membranes were blocked in 5% skim milk powder in TBSTw (Tris-buffered saline (150mM NaCl, 20mM Tris pH7.5, 0.05% Tween-20) for 1h and incubated for 16h with primary antibody in TBSTw. After washing three times 5min with TBSTw, membranes were incubated with HRP-conjugated secondary antibody (1:10,000 BioRad) for 1h at 24°C. Membranes were washed as above and HRP detected by enhanced chemiluminescence (Amersham). Primary antibodies were: rabbit anti-Thiolase (Bodnar and Rachubinski, 1990) mouse anti-Tubulin (Sigma-Aldrich 1:1000).

### Human *PEX7* and Drosophila *Pex7* cDNA cloning and transfection

The open reading frame of human *PEX7* cDNA (Braverman et al., 1997) was cloned into pENTR/D (ThermoFisher) using hPex7-Forward (5’-CACCATGAGTGCGGTGTGCGGTGG) and hPex7-Reverse (5’-GGTCAAGCAGGAATAGTAAGACAAG) primers. The *Drosophila* Pex7 cDNA clone was described previously (Baron et al., 2016). Both were transferred into pT-Rex-DEST30 vector using LR Clonase (ThermoFisher). These were transiently transfected into immortalized human fibroblasts using the Amaxa Human Dermal Fibroblast Nucleofector Kit (Lonza).

### Microscopy

Human fibroblasts were fixed for 30 min in 4% paraformaldehyde in phosphate buffered saline (PBS), rinsed twice in PBST (PBS+0.1% TritonX-100) and blocked for 1h in 5% NGS (Sigma) before incubation for 16h at 4°C with primary antibodies. Following 4 washes in PBST, each were incubated in secondary antibody for 16h at 4°C. After 4 washes in PBST, coverslips were mounted using Prolong-Gold (ThermoFisher). Images were captured using a C9100 camera (Hamamatsu) at 130μm vertical spacing using a 100X oil immersion objective (NA=1.4) on a Zeiss AxioObserverM1 microscope coupled to an ERS spinning disk confocal (PerkinElmer). Primary antibodies included: anti-mouse PMP70 (Sigma-Aldrich) (Imanaka et al., 2000); Activated Caspase3 (559565, BD Pharmagen), anti-phosphohistone H3 (Upstate Biotechnology), rabbit anti-SKL (Szilard et al., 1995) and anti-rat Thiolase (Bodnar and Rachubinski, 1990). Secondary antibodies were: AlexaFluor568 donkey anti-mouse, AlexaFluor488 donkey anti-rat or AlexaFluor647 donkey anti-rabbit (Abcam, 1:1000).

Embryos were collected every 16h at 18°C and processed as reported previously (Parsons and Foley, 2013). Antibodies to Futch (22C10) raised by Seymour Benzer, California Institute of Technology, Even-skipped (2B8) and Repo (8D12) raised by Corey Goodman, University of California, were from the Developmental Studies Hybridoma Bank. Anti-Myosin II was from Abcam (ab51098). All primary antibodies were diluted 1:20, and AlexaFluor568 donkey anti-mouse was the secondary antibody (1:1000, Abcam). Embryo TUNEL staining was performed as described previously (Parsons and Foley, 2013).

## SUPPLEMENTARY FIGURE LEGENDS

**Fig. S1.**
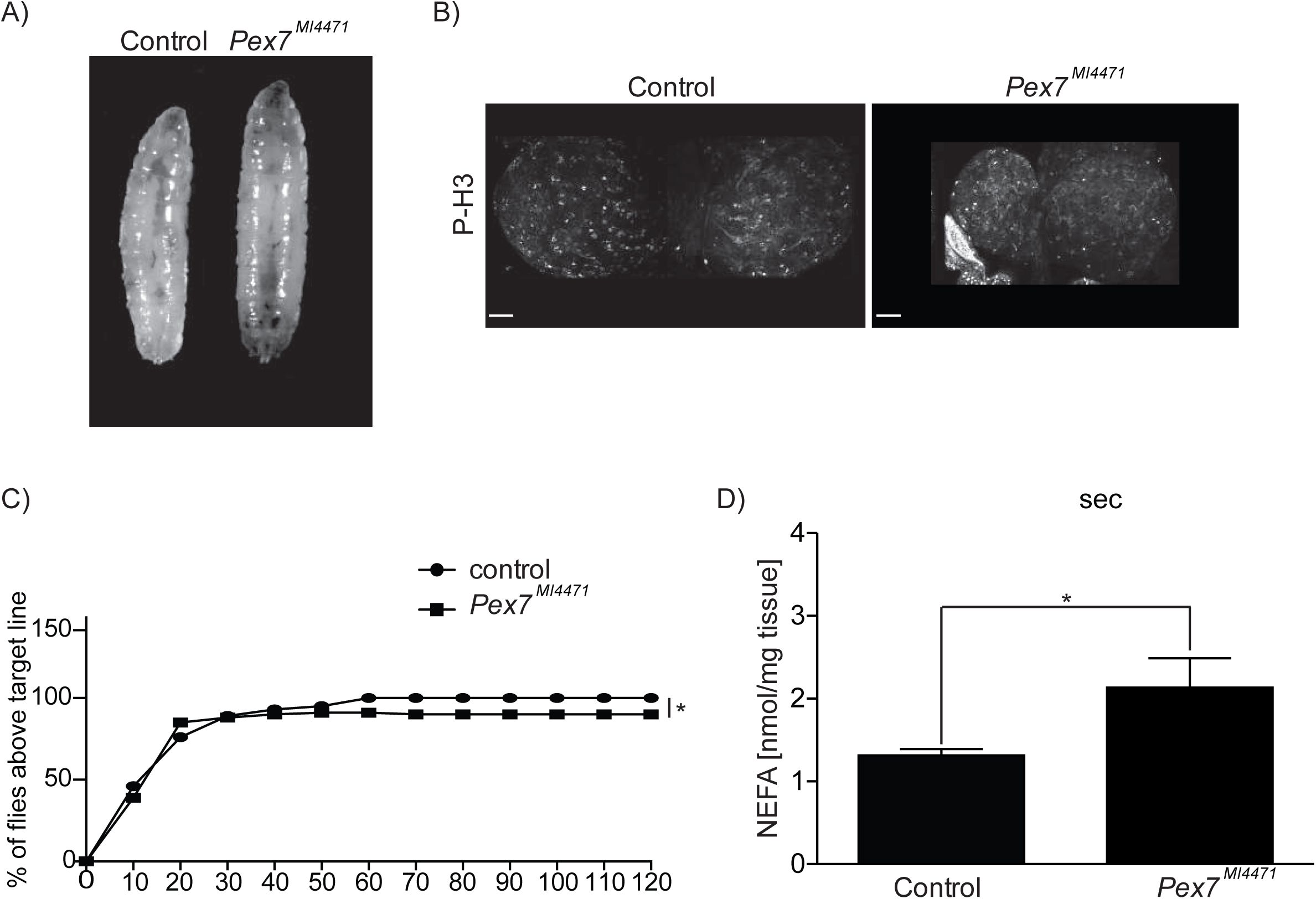
Pex7 mutant show neural defects. (A) Age matched *Pex7^MI4471^*/*Pex7^MI4471^* homozygous larvae are larger than control larvae at the L2-L3 molt. (B) Brains from *Pex7^MI4471^*/*Pex7^MI4471^* larvae exhibit comparable numbers of PH3 positive dividing cells to control flies. Scale bar=1μm (C) The percentage of *Pex7^MI4471^*/*Pex7^MI4471^* flies having passed the threshold line in the climbing assay represented every 10sec is less than control. Significance was determined using Kolmogorov-Smirnov test to compare the distributions of the mutant group to the control. N=12 per time point; * p < 0.05. (D) Amounts of NEFAs in larvae from control and *Pex7^MI4471^*/*Pex7^MI4471^* mutant L3 larvae. Values represent averages of 3 independent experiments ± SD. Significance was determined using Student’s *t*-test; * p < 0.05.

**Fig. S2.**
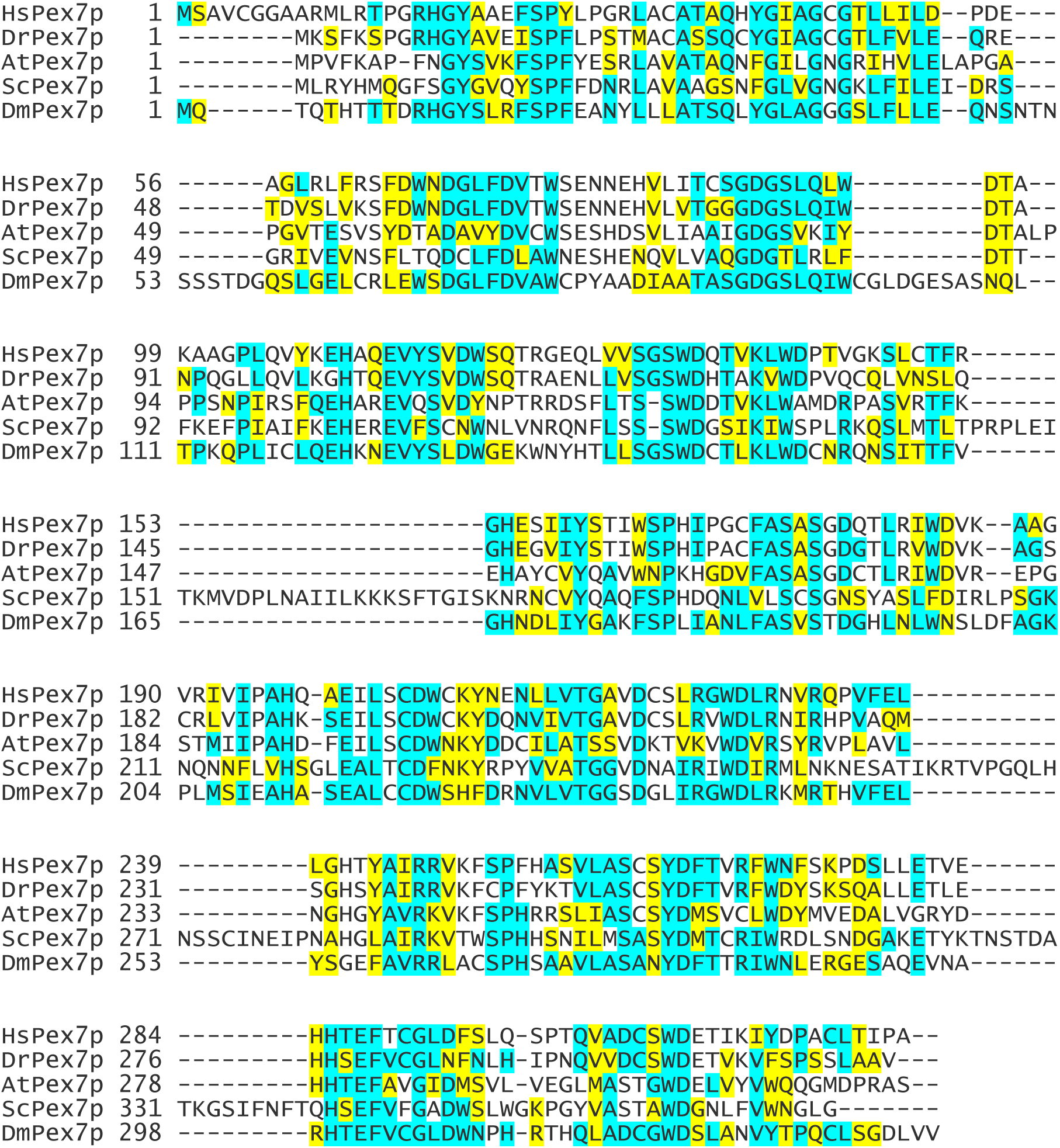
Sequence alignment of Pex7 protein homologues. Sequence alignment of human (HsPex7p, CAG46851.1)), zebrafish (DrPex7p, AAI64338.1), plant (AtPex7p, OAP16388.1), yeast (ScPex7p, KZV12380.1) and fruit fly (DmPex7p, NP_001137914.1). Amino acid sequences were aligned with use of the Kalign program (https://www.ebi.ac.uk/Tools/msa/kalign). Residues that are identical (blue) or similar (yellow) to residues in Drosophila Pex7 are shaded. Similarity rules: G = A = S; A = V; V = I = L = M; I = L = M = F = Y = W; K = R = H; D = E = Q = N; and S = T = Q = N. Dashes represent gaps. Sequence accession numbers are given between brackets.

**Fig. S3.**
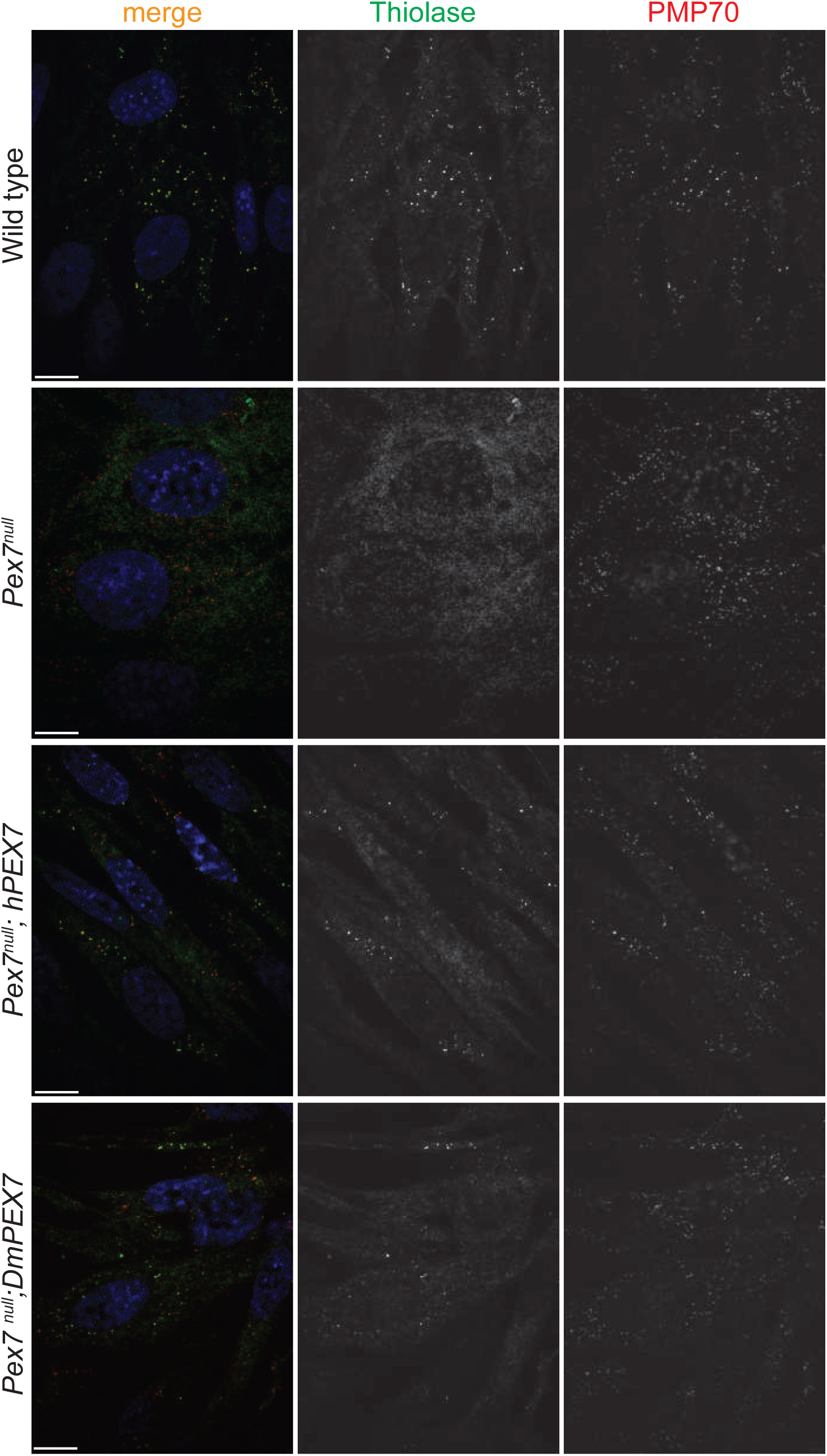

